# The e-Music Box *Roma* : an open research tool for accessible joint music making

**DOI:** 10.64898/2026.07.02.736121

**Authors:** Sara F. Abalde, Félix Bigand, Lorenzo Orciari, Claudio Lorini, Peter E. Keller, Alberto Parmiggiani, Marco Crepaldi, Giacomo Novembre

## Abstract

Joint music making offers an ecologically powerful framework for investigating human social interaction and synchronization. Yet, experimental paradigms often rely on traditional instruments that limit accessibility, reproducibility, and experimental control. In parallel, the use of music for therapy and rehabilitation is expanding, motivating the development of digital musical instruments that can serve research, educational, and clinical purposes. Here, we introduce the e-Music Box Roma (eMB Roma), an open, reproducible digital musical instrument designed to study music making behavior regardless of musical training. The eMB Roma plays preregistered music with tempo controlled by hand rotary movements. Building on the original e-Music Box (Novembre et al., 2015), the eMB Roma retains its intuitive rotary hand control while introducing major innovations: a fully open and 3D-printable design, modular hardware with integrated slider and button controls, polyphonic output with multiple simultaneous instruments, and MIDI compatibility. Additionally, a dedicated graphical user interface allows real-time monitoring, experiment control, device synchronization (like neuroimaging or motion capture devices), and both solo and joint music-making paradigms. The eMB Roma provides a flexible and accessible platform for research contexts, allowing experimental control, reproducibility, and future extensions. Its open design and modularity make it suitable not only for research but also for therapeutic, rehabilitation, and educational applications, where it can support personalized interventions and quantitative assessment of motor performance.

## 1 Introduction

Joint music making provides an ideal model for ecologically valid yet experimentally controlled investigation of human coordination and social interaction (D’Ausilio et al., 2015). However, many musical paradigms used in experimental psychology and neuroscience still rely on traditional instruments, despite their contribution to important conceptual advances (Abalde et al., 2024). This reliance limits accessibility, reproducibility, and experimental control. At the same time, music is gaining increasing attention for its therapeutic and rehabilitative potential (Altenmüller & Schlaug, 2015; Kavety, 2019; Partesotti et al., 2025; Sihvonen et al., 2017; Thaut & McIntosh, 2014). This convergence of interests has motivated the development of Digital Musical Instruments (DMIs), technological systems that enable flexible mappings between human actions and sound production (Miranda & Wanderley, 2006). DMIs can have diverse applications, including research, education, and clinical contexts (Duarte et al., 2023; Feng et al., 2024; Jack et al., 2020; Lindetorp et al., 2023; Zayas-Garin & McPherson, 2022).

In addition, research on human predispositions for music has traditionally prioritized music perception over production, since performance-based paradigms often require prior musical training and are thus less accessible to näıve participants. As a consequence, empirical music research has largely concentrated on auditory processing for much of the twentieth century, with musical movement and action receiving comparatively little systematic attention until more recent decades (Keller & Rieger, 2009; Toiviainen & Keller, 2010). This emphasis on perceptual approaches within music research may, in turn, reflect a broader historical tendency in experimental psychology and cognitive neuroscience to overlook the systematic study of motor control behavior (Rosenbaum, 2005). Consequently, there has been a growing interest in developing accessible, open, affordable, and scientifically validated DMIs that bridge artistic expressivity and behavioral measurement. A variety of musical interfaces have been proposed for this purpose, ranging from motion-capture–based systems (Maes et al., 2024; Wanderley & Depalle, 2004) to interfaces focused specifically on tapping movements (Coorevits et al., 2020; Fink et al., 2022). These systems generally consist of gestural controllers that drive the musical parameters of a sound synthesizer in real time (Miranda & Wanderley, 2006).

Among these instruments is the original *e-Music Box* (eMB): a digital instrument that translates hand rotary movements into musical playback in real time (Novembre et al., 2015). Users rotate a handle on the eMB device to generate musical output, controlling the tempo of preloaded musical sequences, which are played note by note based on rotation angles measured from the hand movements. This technology provided a simplified yet musically expressive interface that allowed participants with or without musical training to generate melodies through hand rotation movements (Novembre et al., 2015). Critically, it enabled controlled investigations of music making by not only allowing participants to generate preregistered musical sequences through hand rotary movements, but also by capturing fine-grained kinematic data that enabled subsequent quantitative analyses, both linear and nonlinear (Pikovsky et al., 2001; Repp & Su, 2013). The instrument has since been used in several studies of musical joint action, and even adapted for music therapy purposes, across multiple research groups (e.g., Liebermann-Jordanidis et al., 2021; Nicol et al., 2024; Novembre et al., 2019; Zhou et al., 2023). Importantly, reports from therapeutic applications involving older adults, including people with dementia, also highlighted positive user experiences, including enjoyment of the instrument’s creative and collaborative aspects, satisfaction associated with successful synchronization, and appreciation of familiar and aesthetically pleasing musical outcomes, with smiles, laughter, singing, and spontaneous movement providing further evidence of engagement and enjoyment (Nicol et al., 2024)..

Despite its impact, the first-generation eMB had practical limitations that motivated the development of the e-Music Box Roma (eMB Roma). The new design preserves the original mapping of rotary motion to discrete musical events while introducing targeted improvements to address key shortcomings: (i) a robust, 3D-printed enclosure and assembly process fully open and reproducible; (ii) native support for reproducing polyphonic musical sequences based on the Musical Instrument Digital Interface (MIDI), with multiple instrument streams enabling simultaneous notes and richer textures; (iii) an integrated workflow for running experiments, loading musical material, and logging performance data; and (iv) absolute angular readout for consistent, comparable motion measurements across sessions. These enhancements are summarised here and described in detail below.

First, whereas the construction of the original eMB relied on existing commercial hardware (i.e., DJ Hero controller; see Figure 1A top), the eMB Roma provides a fully open, 3D-printable design with freely available source code and off-the-shelf electronic and mechanical components (illustrated in Figure 1A bottom), representing a key advancement for shareability, reproducibility, and open science. The device was conceived as modular, allowing individual components (e.g., input controls and housing elements) to be independently modified or replaced within the 3D-printed structure to accommodate different experimental needs. In its current implementation, the instrument includes a linear slider and a push button, extending the interaction capabilities beyond those of the original eMB. These additional controls could enable users to manipulate musical parameters such as loudness, pitch, and timbre (e.g., by switching instruments), thereby broadening the scope of potential applications.

**Figure 1:**
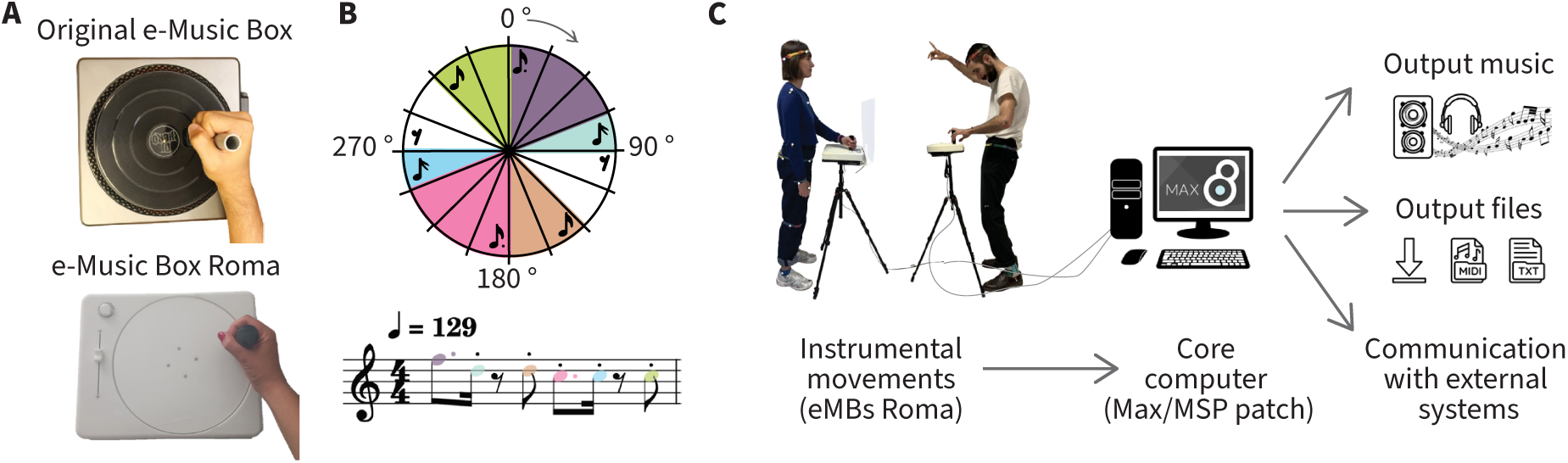
e-Music Box Roma system overview: **A.** Aerial view of the original e-Music Box (Novembre et al., 2015, top) alongside the current version, the e-Music Box Roma (eMB Roma; bottom), shown for visual comparison. **B.** Illustration of an example sixteen-segment discretization of the full 360° rotation (22.5° per segment), mapped onto the sixteenth-note grid of a 4/4 bar. Alternative segmentations and rhythmic mappings can be implemented through the system configuration. **C.** Example of an experimental setup using the eMB Roma system, illustrating the main data flow and outputs.

Second, the eMB Roma supports polyphonic MIDI instruments, enabling the production ofricher musical material beyond monophonic melodies and across a wide range of instrument types, from keyboards to percussion. General MIDI instruments can be dynamically assigned, and multiple notes can be triggered simultaneously, whereas the original eMB generated musical output by sequentially sampling monophonic sounds from a set of pre-rendered WAV files.

Third, the eMB Roma consolidates experimental control, real-time monitoring, and device synchronization into a single graphical interface implemented in Max 8 (Cycling ‘74), which serves as a central hub for controlling multiple eMB Roma devices and, when needed, additional recording systems (e.g., EEG or motion capture; see Figure 1C). This directly addresses two practical gaps of the original system. First, the original implementation depended on the commercial Presentation software (Neurobehavioral Inc.) to translate rotation data into sound, requiring per-computer licenses and the integration of separate software environments for experimental control and stimulus generation. The second gap was the lack of an integrated and operator-editable control layer for running experiments and routing data to external systems. By replacing the Presentation dependency with a Max/MSP patch, the eMB Roma consolidates these functionalities within a single environment. Moreover, although both platforms require licenses, Max offers a lower-cost perpetual licensing model, and compiled patches can be executed across multiple machines without additional licensing fees, with licenses required only for editing. The system also provides a unified graphical user interface (GUI) that operators can adapt without outsourcing basic customizations. Together, these changes reduce technical barriers to deployment and make the platform easier for laboratories to adopt and extend.

Finally, the eMB Roma enhances precision acquisition by using a digital magnetic encoder that registers the absolute rotational phase of each timestamp, ensuring that identical angular positions correspond to identical musical outputs across trials. This constitutes an improvement over the original eMB, which relied on angular displacement (i.e., 45° per quaver note) relative to an arbitrary reference point (taken from the position of the handle when switching on the device). As a result, in the original system, the angular data depended on the initial handle position at startup, potentially introducing inconsistencies in the mapping between movement and sound across trials and devices. In the eMB Roma, rotary hand movements are continuously captured by an embedded sensor, converted into absolute angular values, and transmitted to the core computer that controls the presentation of musical stimuli and data acquisition. By default, the full 360° rotation range is discretized into sixteen equal segments (22.5° each), corresponding to a musical grid quantized to sixteenth notes in a single bar of 4/4 meter, so the rotation angle triggers the note assigned to the current segment (Figure 1B). We adopt this 4/4 sixteenth-note mapping as a convenient default for many Western experimental paradigms (Savage et al., 2015), but the same approach generalises to other meters and subdivision schemes: operators can change the number of angular segments in the configuration files to implement triplet subdivisions (e.g., 12 segments for 4/4 triplets), compound meters (e.g., 12 segments for 6/8 at sixteenth resolution), or irregular meters (e.g., 7 or 14 segments for 7/8 at eighth or sixteenth resolution). This movement-to-note mapping can be customized via the configuration files, allowing operators to define alternative rhythmic structures or resolution levels (see section 3).

By integrating these features, the eMB Roma follows an open and modular design, providing an openly available tool that can support reproducible research on musical interaction, motor coordination, and timing. Its architecture enables flexible adaptation to diverse experimental paradigms, ranging from single-participant tempo-control tasks to multi-user studies of synchronization and cooperation. Beyond research applications, the eMB Roma also holds potential for clinical and therapeutic interventions by offering an engaging movement-based task that produces music, either individually or jointly across multiple instruments, while simultaneously recording high-resolution performance data. These features may support ther-apeutic and clinical research by providing clinicians with objective trial-level measures that could prove useful for assessment and longitudinal monitoring. The required rotary movement is accessible for individuals with a range of mild to moderate motor impairments, while reducing movement-related artifacts during simultaneous brain imaging techniques such as electroencephalography (EEG), compared with tasks involving vocalization or larger body movements. All hardware designs, firmware, and control software to build the eMB Roma are freely available through the eMB Roma repository, fostering transparency, cumulative methodological development, and future extensions.

In the following sections, we describe the design rationale and technical implementation of the eMB Roma system. For simplicity, throughout the remainder of the manuscript we use the abbreviation *eMB* to refer interchangeably to the present eMB Roma system unless otherwise specified. We use the term *operator* to denote the person running the eMB system and managing the GUI (e.g., researcher, experimenter, clinician, or other system controller), and *user* to denote the person interacting with the eMB (e.g., experimental participant, patient, or artist; essentially any performer). We first present the hardware architecture, detailing how to build the eMB and providing sufficient technical information to support replication and modification by operators.We then describe the software architecture underlying the eMB, explaining both the logic and use of the GUI in order to run different experimental paradigms and adapt the system to specific operator needs. Finally, we discuss the configuration dimensions of the eMB Roma system, outline potential fields of application and use cases in domains such as music cognition, sensorimotor coordination, social interaction, and clinical and therapeutic research, describe the system’s open and extensible design, and conclude by addressing future developments, current limitations, and broader perspectives for the eMB Roma.

## 2 Building the eMBs: Hardware design and construction

### 2.1 Mechanical design and fabrication

Each eMB Roma unit is fabricated using custom 3D models provided as openly available STL files. The fully assembled enclosure (Figure 2A) measures approximately 33 × 27 × 5 cm, and encloses all mechanical and electronic components. The fabrication process comprises three main structural groups: the rotary disk group, the cover group, and the base group, as illustrated in Figure 2B, from top to bottom.

**Figure 2:**
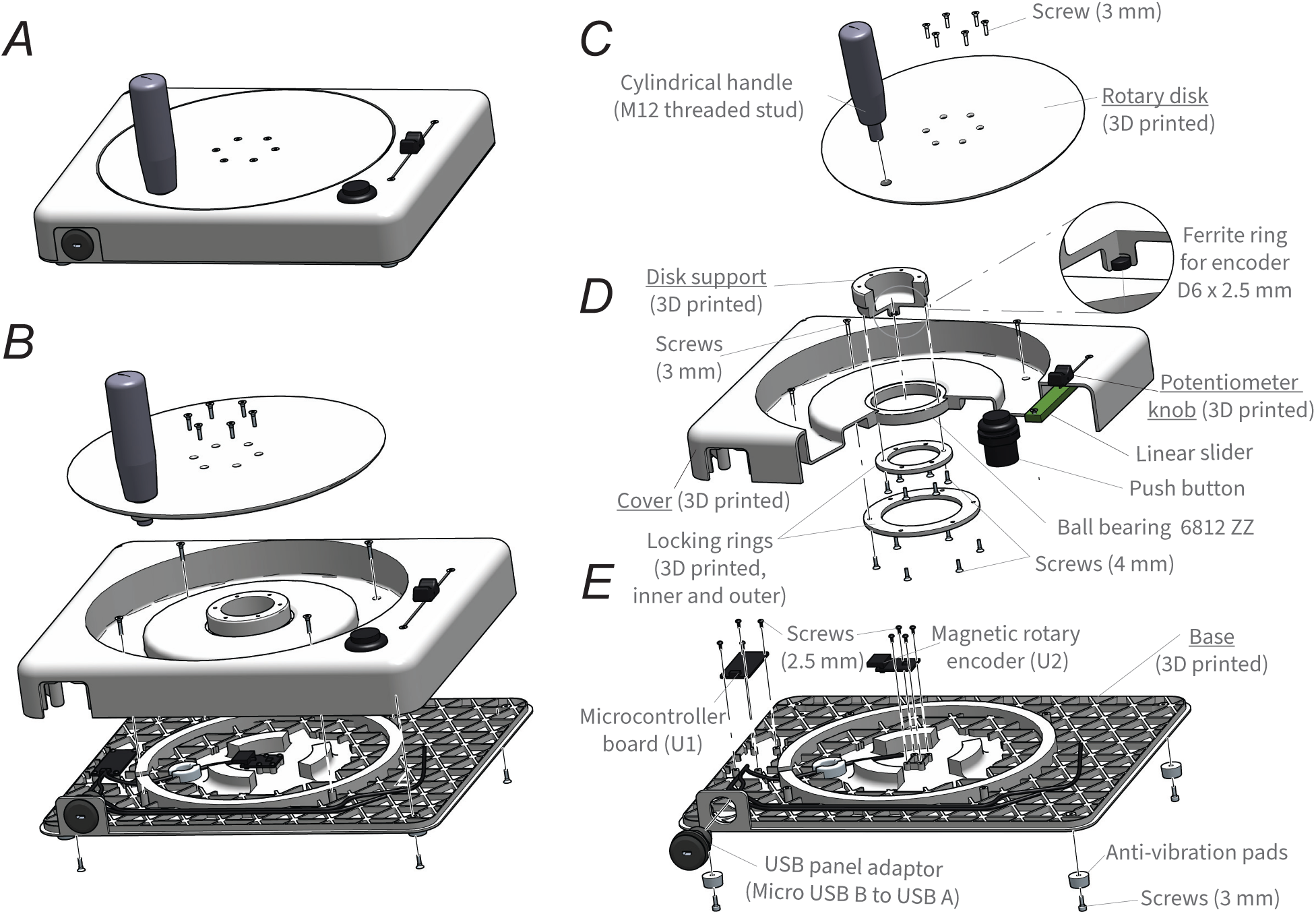
Mechanical design of the eMB Roma. **A.** Fully assembled device with closed enclosure. **B.** Main structural groups (rotary disk, cover and base; 3D printed) shown separated and without internal components. **C-E.** Exploded views of each group, detailing all mechanical and electronic elements: **C.** rotary disk, **D.**cover, and **E.**base group. Key components include the rotating cylindrical handle, ball bearing (6812 ZZ), locking rings, disk support, linear slider, push button, microcontroller board, and magnetic rotary encoder (the latter three are described in the Electronic Components section). Underlined components are 3D printed in Nylon PA 6-6. All screws used in the assembly are self-tapping, with diameters of 2.5 mm, 3 mm, 4 mm, corresponding to standard M2–M4 sizes.

All 3D printed parts were produced in Nylon PA 6-6, selected for its rigidity and durability. However, the open-source nature of the design allows for alternative materials to be used if required by specific experimental constraints. While most components are either 3D printed or electronic (see section 2.2), additional mechanical elements include the handle, the ball bearing, and the ferrite ring.

Assembly is primarily achieved using M3 (3 mm) ultra-flat hex socket head self-tapping screws, which provide a low-profile finish and mechanical robustness. Other screw sizes (e.g., M2 and M4), also self-tapping, are used where indicated in Figure 2C-E.

The base group (Figure 2E) forms the structural foundation of the device. It includes mounting holes for four integrated feet, one at each corner, providing stability and anti-slip. The base also features dedicated placeholders for the microcontroller board and the magnetic rotary encoder (see section 2.2), allowing secure attachment without interfering with device functionality. It also incorporates a lateral placeholder to accommodate the USB connector for straightforward external access.

The base group (Figure 2E) forms the structural foundation of the device. It includes mounting holes for four integrated feet, one at each corner, providing stability and anti-slip support. Different feet can also be selected to accommodate different installation requirements, such as fixation to external surfaces or experimental tables. The base further features dedicated placeholders for the microcontroller board and the magnetic rotary encoder (see

section 2.2), allowing secure attachment without interfering with device functionality. It also incorporates a lateral placeholder to accommodate the USB connector for straightforward external access.

The cover group (Figure 2D) is designed to enclose the base while supporting the rotary disk assembly. Central to this group is a thin-section ball bearing (6812 ZZ; 60 × 78 × 10 mm, bearing steel, double metal shield), which ensures smooth and precise rotation. Axial alignment is maintained using three custom 3D-printed components: two locking rings (one smaller, one larger) and a disk support. The locking rings secure the bearing to the cover, while the disk support attaches to the rotary disk and houses a ferrite ring core positioned above the magnetic rotary sensor (see section 2.2 and zoomed region in Figure 2D). The inner race of the bearing is fixed to the rotary disk, and the outer race is anchored to the enclosure, enabling stable rotation directly above the sensor. The cover of the eMB also includes the placeholders for the linear slider and the push button (see section 2.2).

The rotary disk (Figure 2C), which enables user-controlled rotational input, is 3D printed and designed to interface with a rotating cylindrical handle equipped with an M12 threaded stud. This threaded stud fits directly into the central mounting point of the disk, allowing secure attachment. In our implementation of the eMB Roma, we selected a handle made of phenolic resin with a stainless-steel threaded shaft (total length = 124.5 mm; Art. 24530.0271). However, the open-source design permits the use of alternative handles, provided that they include a compatible threaded shaft or that the rotary disk model is modified accordingly. To improve mechanical stability during repeated rotational movements, the handle may optionally be secured using a thread-locking compound after assembly. The rotary disk is then attached to the disk support, completing the mechanical assembly of the eMB Roma.

### 2.2 Electronic architecture

Each eMB device integrates a rotary magnetic encoder, a linear slider, and a push button, all interfaced to a single microcontroller responsible for data acquisition and USB communication with the core computer. These components and the associated mounting hardware are illustrated individually in Figure 3A, and shown in example photographs in Figure 3C. All peripherals shared a common ground reference with the microcontroller, and all digital and analog signals operated at 3.3 V logic levels.

**Figure 3:**
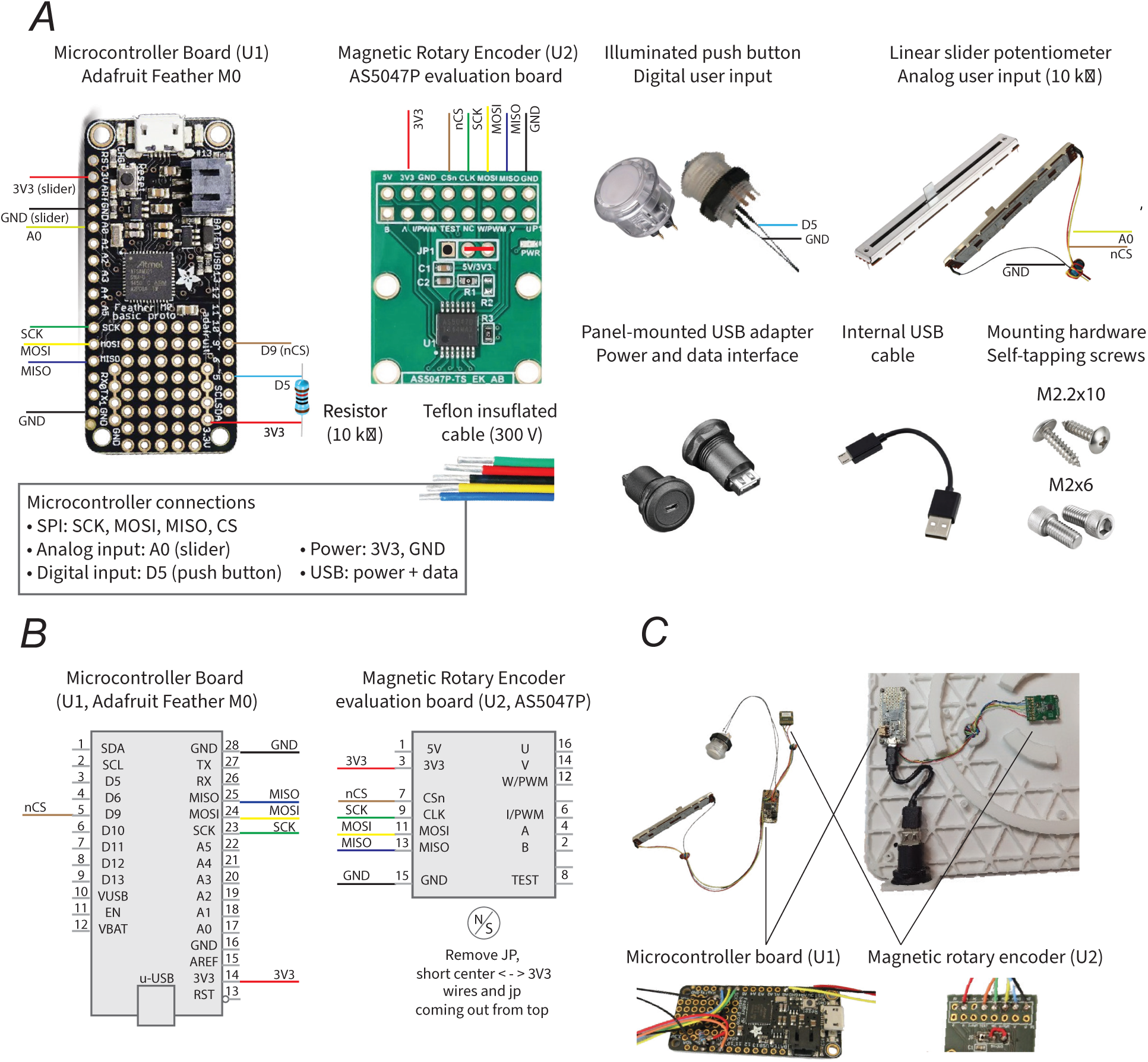
Electronic architecture. **A.** Components used to implement the electronic setup with connections, including an illuminated push button, a linear slider potentiometer, a panel-mounted USB adapter with internal USB cabling for power and data communication, and mounting hardware. **B.** Connection diagram showing the electrical interfaces between the microcontroller board (U1, Adafruit Feather M0) and the magnetic rotary encoder evaluation board (U2, AS5047P). The encoder communicates with the microcontroller via the Serial Peripheral Interface (SPI) and is powered at 3.3 V. **C.** Photographs illustrating the internal wiring and physical integration of the electronic components. Left: cabling connections between the microcontroller board and the user input devices (push button, linear slider) and the magnetic rotary encoder. Right: assembled electronics mounted to the 3D-printed base of the enclosure, showing the microcontroller board, rotary encoder, and panel-mounted USB adapter. The slider and the illuminated push-button are not shown because they are installed in a different location.

The central controller of the eMB is an Adafruit Feather M0 microcontroller (Arduino-compatible, ARM Cortex M0, Atmel ATSAMD21G18). The microcontroller is powered via a USB connection, which supplies 5 V and is regulated onboard to 3.3 V for the microcontroller and all connected peripherals. The same USB connection is used for bidirectional data communication with the core computer. Internally, the microcontroller connects to a panel-mounted Micro-USB-B–to–USB-A adapter (Adafruit 4213, Ø30 mm) using a USB-A–to–Micro-USB-B cable (150 mm), providing a single external interface for both power delivery and data communication (Figure 3A, C).

Rotation of the eMB disk is measured using a magnetic rotary encoder module comprising a magnetic rotary position sensor IC (AS5047, ams OSRAM) mounted on a dedicated evaluation board (EVB_AS5047P). The sensor provides a 14-bit core angular resolution with dynamic angle compensation. The encoder module is mounted within the lower enclosure, directly beneath the bearing structure coupled to the rotary disk (Figure 2D). A ferrite ring core (FL1, 1760 nH, 2914 package, 6 × 4 mm, 2/3 turns) is integrated into the magnetic assembly to improve magnetic field stability and reduce susceptibility to external interference, as illustrated in the zoomed region of Figure 2D.

To ensure compatibility with the 3.3 V logic levels of the microcontroller, the AS5047P evaluation board was configured by modifying jumper JP1, as shown in Figure 3A. Specifically, the default jumper connection was removed, and the center pad was solder-bridged to the 3.3 V pad, thereby selecting 3.3 V supply and logic operation in accordance with the manufacturer’s specifications. The encoder evaluation board includes local power conditioning and signal-filtering components; all passive elements required for encoder operation (including decoupling capacitors and configuration jumpers) are located on the encoder printed circuit board (PCB), and no additional external passives were required. The encoder communicates with the microcontroller via a Serial Peripheral Interface (SPI), with the specific signal connections between the microcontroller board and the evaluation board shown in the connection diagram of Figure 3B.

In addition to the rotary input, each eMB includes a linear slider implemented using a 10 kΩ PTB0143 linear potentiometer (100 mm travel, 20% tolerance), fitted with a 10 mm knob, and an illuminated arcade-style push button with integrated LED (Adafruit), mounted as shown in Figure 3A and C. The slider position is read via an analog input of the microcontroller. The push button is read through a digital input and connected using an external 10 kΩ resistor, as shown in Figure 3A, which provides a defined logic level when the button is not actuated and prevents a floating input state. These additional user inputs are sampled alongside the rotary encoder data and transmitted to the core computer over the USB interface. All electronic components were powered exclusively via the USB connection, and no external power supplies were required.

### 2.3 Connecting the eMB to the core computer: firmware and data communication

Each eMB operates using custom firmware that is uploaded to the microcontroller once during initial device setup and does not require modification during routine use. The firmware is developed in the Arduino environment and is publicly available in the eMB repository (https://github.com/sarafernandezab/eMusic Box Roma). Communication with the magnetic rotary encoder is implemented using the AS5047P Arduino library (https://github.com/jonas-merkle/AS5047P), which must be installed as a dependency.

The firmware reads the rotary encoder via the Serial Peripheral Interface (SPI) at a bus speed of 100 kHz and continuously acquires the rotation angle, linear slider position, and push button state. Sensor data are streamed in real time to the core computer via a serial (COM) port at a baud rate of 9600 bps using ASCII-encoded values in a fixed-order message format.

Because fine mechanical alignment of the encoder magnet cannot be guaranteed during assembly, device-specific angular offsets are corrected in software. After assembling each eMB, the device is connected to the computer, and raw angle data are sent via the serial monitor of the Arduino development environment. Calibration is performed by modifying a single firmware parameter (value_offset_zero), which defines the angular value corresponding to the desired zero position by subtracting the offset from the encoder readout and wrapping the result within the 0–360° range (i.e., (readAngleDegree() - value_offset_zero) % 360). The modulo operation ensures that angular values remain within a 0–360°range. This calibration procedure is performed once per device after assembly and does not need to be repeated unless the encoder or magnet position is modified. Calibrating this parameter ensures consistent angular alignment across devices while allowing alternative zero-reference positions when required.

The direction of rotation (clockwise or counterclockwise) can be configured in the firmware. By default, the eMB is set to clockwise rotation, but this behavior can be inverted by changing the rotation_direction parameter from CW to CCW. Internally, the firmware first aligns the raw encoder readout to the calibrated 0–360° range and, when counterclockwise behavior is selected, mirrors this value before streaming it to the computer. After changing the rotation direction, the zero-reference parameter (value_offset_zero) may need to be verified or adjusted to preserve correct angular alignment.

Once calibration and any firmware modifications are completed and the firmware is uploaded, the Arduino development environment must be closed so that the serial port becomes available to the Max/MSP control patch. During experiments, serial communication is handled by a custom Max/MSP patch (Max 8, Cycling ‘74), which manages data reception, experimental control, and synchronization across multiple eMB devices and external systems, as described in the following section.

## 3 Playing the eMBs: Software architecture and experimental control

A dedicated GUI implemented in Max/MSP provides a unified framework for interactive musical stimulus presentation and experimental control of the eMB system. The interface allows operators to load configuration files, select musical stimuli and associated instruments, define experimental parameters, monitor participants’ performance in real time, acquire eMB data streams, and transmit synchronization triggers to external recording systems.

### 3.1 Configuration and initialization

At the start of each session, two configuration files are loaded into the Max patch: a global configuration file and an experiment configuration file (with contents detailed below; illustrated in the violet block in Figure 4). Both are plain-text documents (.txt) that can be automatically generated from a custom Python script openly available, which can be easily modified to suit the operator’s needs. The global configuration file contains the general settings for the experimental setup, such as the eMB ports, sampling rate, file paths (both for output and input files), and which instruments are assigned to each channel. The experiment configuration file, specific for each experiment, indicates what the specific parameters to be set for each trial are, e.g., song, instruments, or the condition of that trial. This architecture allows experiment-specific randomization and reproducible session setup. Upon initialization, the Max patch automatically reads the global configuration file and establishes serial connections with each eMB (illustrated within the violet block in the diagram of Figure 4). The GUI then displays the loaded configuration, as well as the trial-specific configuration if applicable, allowing visual verification before beginning data acquisition.

**Figure 4:**
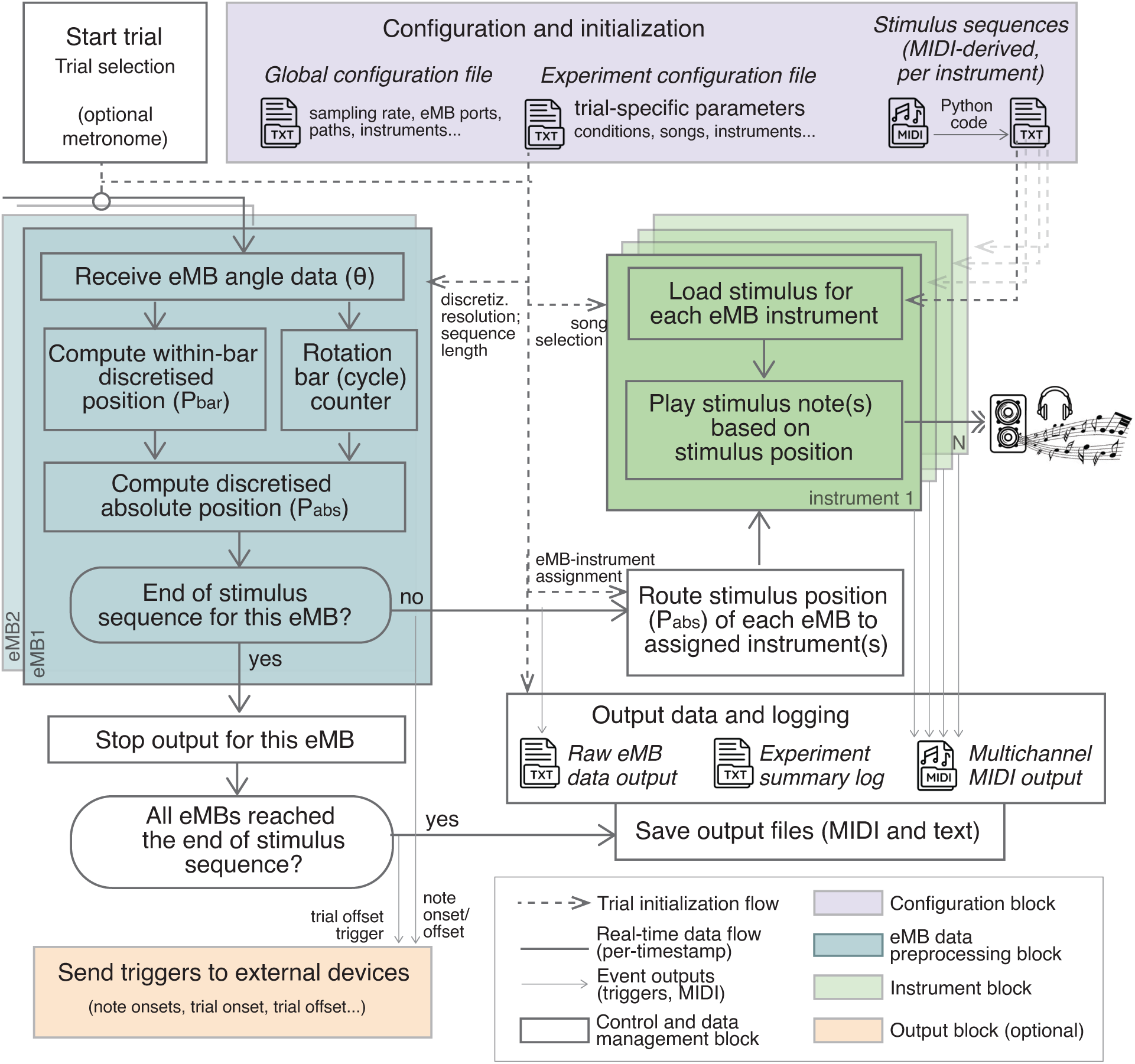
Diagram of the main data flow in the software architecture implemented in Max/MSP. The top (violet) block represents files loaded during patch initialization, including a global configuration file, a trial-specific configuration file, and musical stimulus files. The blue block illustrates the real-time data processing performed independently for each eMB. When a trial starts, incoming rotary angle data from the eMB(s) are decomposed into a rotation cycle count and a within-bar phase (default discretization: 16 segments per rotation, corresponding to the sixteenth-note structure of a 4/4 bar). The green block represents stimulus handling, where discretized eMB positions are mapped onto musical events and routed to the assigned instrument(s) for each eMB. When a discretized position corresponds to a note defined in the stimulus file (previously loaded for the corresponding instrument), the corresponding event is generated and written to the multichannel MIDI output. Optional triggers to external devices can be emitted at predefined points in the data flow (orange block) and logged alongside experimental data. Trial execution continues until all eMBs reach the final discretized position of the stimulus sequence, after which trial-specific output files (MIDI and text-based logs) are generated and saved.

The global configuration file contains parameters common across experiments that can be optionally changed, including the sampling rate (by default 200 Hz), the assignment of eMB (COM) ports, serial communication parameters, paths of loaded stimulus sequences (MIDI-derived songs), output files, and other optional parameters such as metronome settings (pulses number, pitch and duration), and external connection ports to send trigger outputs signals to measure devices, such as EEG.

The experiment configuration file defines the trial-specific parameters for a given experiment, in a trial-by-trial manner. Each row in this file represents one trial and it can include the following columns: (1) trial number, (2) trial condition, (3) song name, (4) song number, (5) number of bars of the song, (6) eMB1 instrument assignation, (7) eMB2 instrument assignation (if applicable), (8) metronome (bpm), (9) number of metronome beats to be played at the start of the trial. Song name and song number represent the name of the file containing the MIDI-derived stimulus sequence to be loaded for the respective trial (see next section 3.2), and the number is redundant information to identify the song, avoiding full names within the patch. These trial files can be batch-generated for all experiments via a custom Python script, openly available. This script can be easily adapted to control the randomization of conditions or other experimental requirements like the song or instrument assignment per trial.

### 3.2 Musical stimuli

All musical materials for each instrument stream must be formatted to be interpreted and synchronized precisely with the rotational data from the eMBs, ensuring temporal alignment with participants’ movements and rotational discretization. These sequences of musical stimuli are a simplified text-based representation converted from standard MIDI files of each instrumental part. They should be available in the assigned paths of loaded stimulus sequences for each of the assigned instruments (both set from the global configuration file) at the start of the experiment.

By default, the musical grid is discretized into sixteen equally spaced steps per full rotation of the device, corresponding to the sixteenth-note structure of a 4/4 bar, although this mapping can be adapted to support other discretizations and metrical signatures. Each step represents a fixed temporal frame within which one or more notes may occur, as illustrated in Figure 1B. The text files are encoded following the MIDI protocol, so each note is represented by three values: pitch, velocity, and duration. Velocity represents the intensity of the key press, determining the resulting loudness of the note (the higher, the louder), while duration specifies its temporal length. In polyphonic stimuli, where multiple notes occur within a single frame, each note is represented by a corresponding triplet of pitch, velocity, and duration values within the same row. During playback, the Max patch retrieves the appropriate musical events for each frame and instrument stream in real time by reading each row (e.g., via the coll object) based on the current angular position mapping, and advances through the sequence according to the rotation of the eMB handle (see next section). Frames without musical events are mapped to silence, with all corresponding MIDI values set to zero, thereby preserving the rhythmic structure of the original musical stimulus sequence. All conversions from MIDI to text files are implemented using Python libraries (i.e., Pretty MIDI; Raffel and Ellis (2014)) to facilitate reproducibility, and these conversion scripts are available. Note that this conversion is done for each instrument separately, so in case of having multiple instruments in the same MIDI file, they need to be separated in different files beforehand (with filenames ending with the respective instrument name). The script allows polyphonic MIDI sequences and batch conversion of multiple songs in the same folder and automatically extracts metadata such as tempo and bar count from the MIDI file header, enabling operators to generate compatible stimuli from any MIDI source. The resulting text format is intended to ensure compatibility and enable the generation or modification of the MIDI-derived sequences without specialized programming expertise beyond basic MIDI notation and protocol conventions (MIDI Manufacturers Association, 1996), either by changing the Python code or manually modifying the text file in any text software. For example, to transpose the notes of the original MIDI sequence to a lower octave, the processed pitch value of each note needs to be decreased by 12.

Example ready-to-use musical materials are available as MIDI and .txt files (MIDI-derived songs) in the repository. These are text files converted from eight remakes of famous song refrains from electronic dance music and disco-funk genres used in previous research (Bigand et al., 2024, 2025). They correspond to four different instruments: drums, bass, keyboards, and violin (as the vocal melody), all following a 4/4 meter and spanning 16 bars. In these musical stimuli, we noticed the original percussion and keyboard MIDI sequences had long and almost-overlapping notes, so we reduced their note durations by 20% to prevent excessive overlap between these consecutive notes when rotating the eMB at fast speeds by adjusting the original duration number of each note accordingly (multiplied by 0.2).

Each MIDI-derived text file contains metadata specifying stimuli-specific parameters, such as tempo and the number of note frames, which are automatically extracted from the original MIDI sequences. This metadata can be useful for setting up other parameters in the eMB, such as the stimulus tempo to match the metronome tempo, and the number of frames to define the end of the sequence. The MIDI-derived text files are read by the GUI upon loading, according to the configuration files previously described (also illustrated in the violet block of Figure 4). Individual instrumental parts (for example, melody, bass, or percussion; saved as text files named *<*song*>_<*instrument*>*.txt) are stored in separate files, allowing them to be flexibly assigned, mixed, or synchronised across multiple devices during performance (see section 3.4).

### 3.3 Trial execution and data flow

Each trial is initiated manually by the operator from the GUI. Before starting a trial, the operator selects the desired trial number, which determines the corresponding musical stimuli and configuration settings to be loaded from the trial-specific configuration file. Pressing *Start trial* initializes timing and begins streaming real-time data from each eMB at the specified sampling rate. Optional metronome pulses can be played before the beginning of each trial to facilitate synchronization, according to the parameters defined in the global configuration file (e.g., number of pulses, pitch, and duration).

During data acquisition, the continuous angular position values of each eMB are discretized to match the temporal resolution of the stimulus sequence. While the system supports configurable discretization resolutions, the current implementation maps positions onto a 16-step grid corresponding to the sixteenth-note structure of a 4/4 metrical bar. Thus, each full rotation corresponds to one bar subdivided into *N* equal temporal bins, where in the current implementation *N* = 16. This formulation can be adapted to different temporal resolutions and metrical structures by modifying the discretization parameters and corre-sponding sequence mappings. Accordingly, the discretized position within a bar for each eMB, denoted *P_bar_*, is computed as:

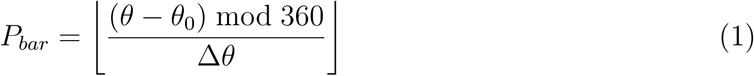

where *θ* is the angular position, *θ*_0_ is the zero-position offset (which should be zero if the device has already been calibrated in the firmware; see section 2.3), and Δ*θ* = 360*/N* corresponds to the angular increment associated with one temporal bin (22.5° in the current 16-bin implementation).

To determine which note in the full song should be played at a given position within an eMB rotation cycle, the system computes the absolute temporal index, denoted *P_abs_*. For example, in the current 16-bin implementation where *N* = 16, the 31^st^ temporal bin—obtained after one complete rotation plus 15 additional bins, corresponds to the 15^th^ position of the second bar. In general, the absolute index is computed by adding the current within-bar position to the number of completed rotations multiplied by the number of bins per rotation:

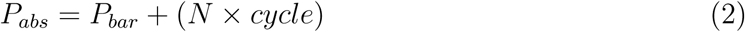

The indices *P_abs_* determine the note to play for each eMB, since each discrete step triggers the corresponding MIDI event within the preloaded instrument file. These operations are illustrated in the soft-blue block of Figure 4, as they are executed independently for each eMB in parallel. Note that if a full-bar metronome is added at the beginning of each song, the musical stimuli should be adapted to have a silent bar at the beginning, and consequently the *P_abs_* should include an initial silent bar. In this case, *P_abs_* should be computed by adding an additional *N* units to account for this offset.

### 3.4 Instrument mapping and interactive musical stimulus presentation

Each eMB can be assigned to one or more MIDI instrument streams (e.g., keyboards, bass, drums, guitar, or multi-instrument combinations), which are pre-defined in the global configuration file according to the general MIDI instrument codes (see Supplementary Tables S1–S11). During a trial, the Max patch dynamically routes each eMB’s absolute position (*P_abs_*) to its assigned instrument channel using a gate structure, as illustrated in the green block of the diagram in Figure 4. This setup enables flexible mapping of devices to instruments across trials, supporting both solo and joint music making paradigms. By default, streams 1–4 correspond to melody piano, bass, drums, and combinations of these instruments, although these assignments can be customized through the global configuration file.

Music generation is implemented by playing preloaded MIDI-derived songs encoded as text files through real-time MIDI streams synchronized with the discrete indices generated by eMB rotation. Each row of the text file contains the note information required to drive the MIDI instruments (e.g., pitch, velocity, duration, and instrument channel). While the user is playing the eMB, the absolute rotational position *P_abs_* is mapped onto these discrete indices, triggering the corresponding note event in the sequence. All instrument outputs are rendered in real time, providing immediate auditory feedback corresponding to the users’ movements.

### 3.5 Data output and logging

Trials end automatically when all the active eMBs reach the end of their respective musical sequences (as illustrated in Figure 4). Then, for each trial, the Max patch produces two output files (an example of a trial with two eMBs is shown in Figure 5):

**Figure 5:**
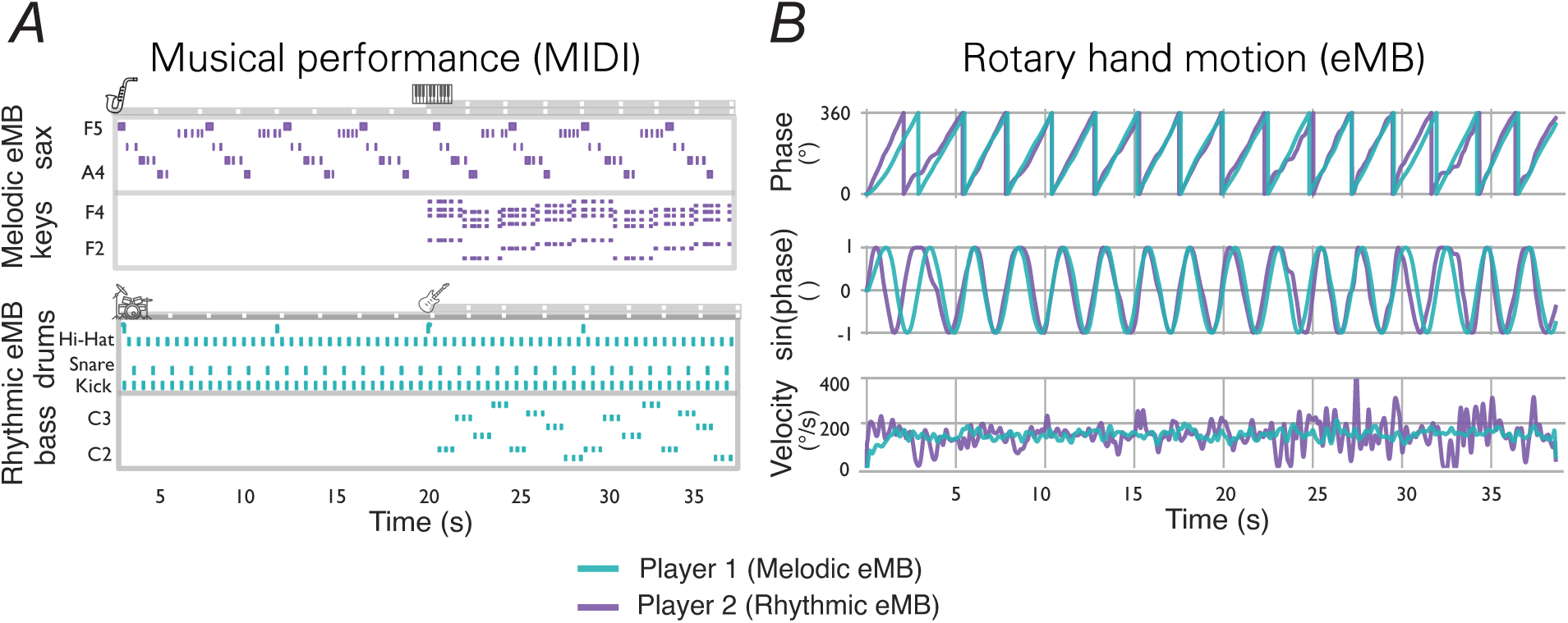
Example of dual music making trial with two eMBs, each controlling two polyphonic MIDI instrumental streams (melodic eMB: saxophone, piano; rhythmic eMB: drums, bass), introduced incrementally. Participants performed an arranged excerpt of “Freed from Desire” using these instrumental streams. **A.** Multitrack MIDI performance data over time (x-axis), with instrument labels shown above each plot; y-axis ticks indicate musical pitch labels for pitched instruments (saxophone, piano and bass) and representative drum classes for percussion (drums). **B.** Instrumental hand rotary motion data from the eMB raw text output files, together with example motion-derived measures. The top row shows the recorded raw phase for each device in degrees; the middle row shows the sine(phase) as a phase-normalized waveform; the bottom row shows the angular velocity in degrees per second.

1. **Raw data text file**, containing per-timestamp values (as rows, at the desired sampling rate) for all connected eMBs within the same file. For each eMB, the following values are stored in separate columns: (1) angular position (*θ*, °), (2) discretized ab solute temporal position (*P_abs_*), (3) completed rotation count (cycle number), and (4) instrument number played (0 = muted; see above). For experiments involving two eMBs, the same four-column structure is repeated for the second device, yielding a total of eight columns. Although the current implementation supports one or two eMBs, the same structure could be extended to additional devices if required.
2. **MIDI file**, containing all musical output streams from the trial, with each instrument in a different channel corresponding to the assigned instruments played by all eMBs.

Additionally, if logging is enabled in the configuration file, a summary log file is also saved. It contains a new row for each executed trial, where at the end of each trial it lists the trial name (of the output trial file), song played, assigned instruments, start and end timestamps, and total elapsed time. The MIDI file is primarily retained for auditory inspection, while behavioral analyses are conducted using the raw data text files (i.e., the angular data logs, which also have higher temporal precision).

### 3.6 Synchronization with external devices

The Max patch optionally facilitates the generation of TTL triggers at specific experimental events (e.g., trial onset, offset, or predefined rhythmic positions) that can be sent through COM ports. These can be set to be sent at predefined rhythmic positions, such as selected note onsets, or more general experimental events, such as trial onset and offset (as illustrated in the orange block in Figure 4). This feature enables temporal synchronization with other recording systems, such as EEG or motion capture, facilitating multimodal acquisition and subsequent data analysis with received triggers.

## 4 System configuration dimensions and example use cases

To illustrate the versatility of the eMB Roma, we describe two primary system configuration dimensions: musical complexity and social configuration. Musical complexity refers to the structure and richness of the musical output that can be produced by eMB Roma. This includes whether the output music is monophonic or polyphonic, which instruments are included, and what musical material these instruments play. The other dimension, social configuration, refers to whether the eMB is played solo by one user or in duet by two users. These two dimensions can be combined and parametrically varied to suit different experimental aims (see Figure 6). Other system configurations that can be relevant for some paradigms are the rotational mapping (rotation direction and starting angular reference) and external device connections. These additional system configuration dimensions, which were not illustrated in Figure 6 for space limitations, can be combined with the main two dimensions to achieve different use cases, as exemplified later.

**Figure 6:**
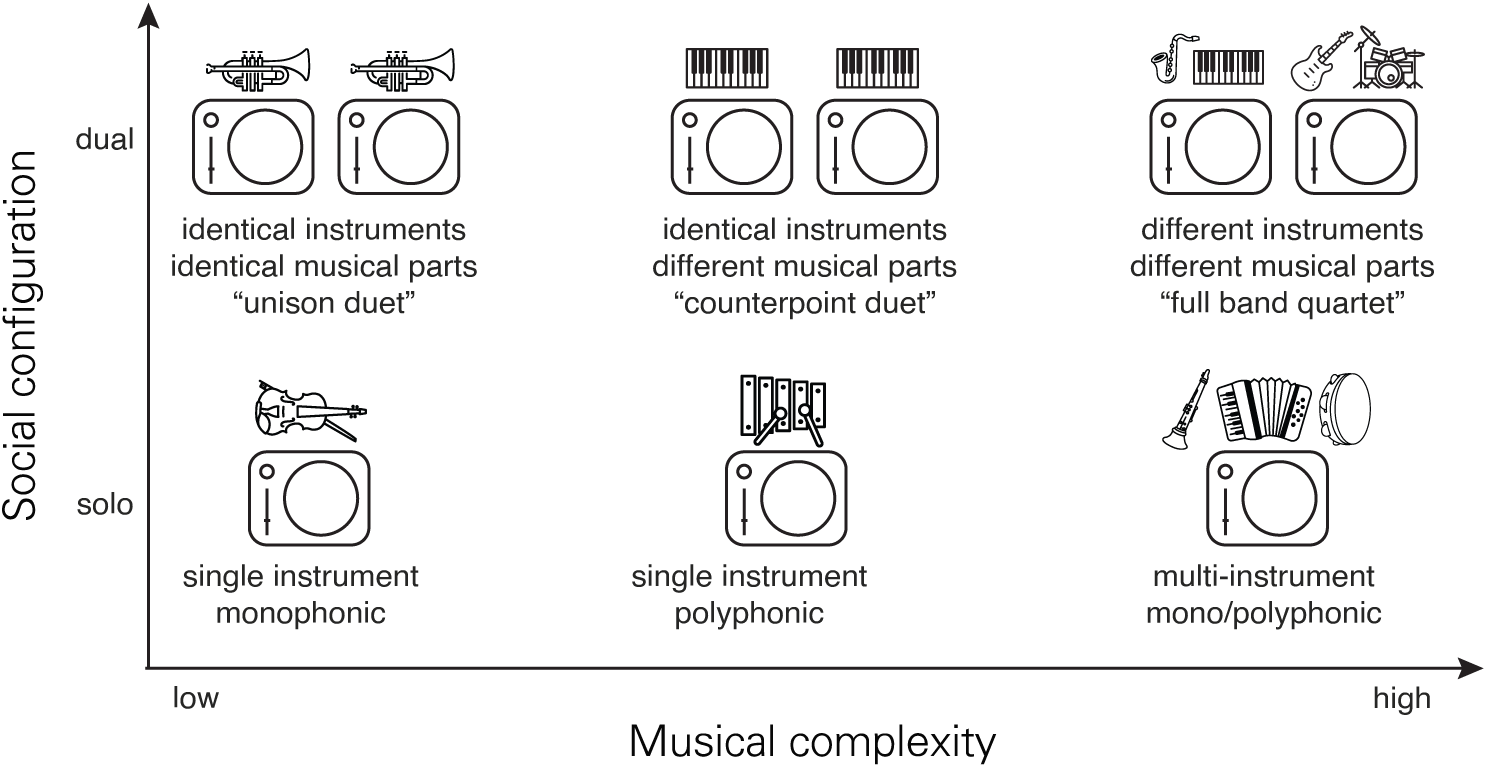
Example use cases plotted across the two configuration dimensions: social configuration (vertical axis: solo vs. dyad) and musical complexity (horizontal axis: low → high). Social configuration is shown for solo and dual setups, but could be extended to multi-user (i.e. including more than 2 users) setups with additional software and synchronization steps. Musical complexity is an abstract continuum from simple (single instrument, monophonic, identical stimuli) to complex (multiple instruments, polyphony, independent stimuli). Example positions on the plot are annotated with illustrative use cases (e.g., “unison duet”, “counterpoint duet”, “full band quartet”) described in the text.

### 4.1 Musical complexity

The musical complexity dimension captures the structure and richness of the musical output produced by each eMB Roma and can be manipulated along several complementary axes: (i) polyphony, (ii) instrumentation, (iii) stimulus relationship, and (iv) real-time parameter modulation. These axes can be combined to create simple, tightly controlled stimuli or rich, multi-stream textures depending on experimental goals, as illustrated along the horizontal axis in Figure 6.

i. **Polyphony.** For each instrument, the eMB reproduces monophonic or polyphonic sequences according to the loaded MIDI-derived stimuli. If a text sequence contains more than one MIDI note at the same timestamp (row), those notes are played simultaneously as a polyphonic chord. Operators can create polyphony for each instrumental stream, either by editing the original MIDI before conversion to text or by modifying the converted text files directly prior to loading them into the Max/MSP patch.
ii. **Instrumentation**. The instruments played by each eMB are set via MIDI program-change codes (see Supplementary Tables S1–S11). The current Max/MSP patch supports up to four independent instrument streams (illustrated as green blocks in Figure 4), although this limit can be extended by adding additional streams in the patch. Each stream receives its own MIDI-derived stimuli, so instrumentation choices determine both timbral variety and the number of parallel musical lines available for manipulation.
iii. **Stimulus relationship across streams**. Because each instrumental stream loads stimuli independently, musical complexity can be configured by choosing whether streams play identical material, complementary/interlocking parts (e.g., melody vs. accompaniment), or independent material. This flexibility supports experiments that probe entrainment to identical signals, coordination on interlocking patterns, or cognitive load when attending to multiple independent streams.
iv. **Real-time modulation via the linear slider**. The linear slider provides continuous control values that are streamed into the Max/MSP patch. Although live mappings (for example, continuous pitch shifts, filter sweeps, or dynamic level control) are not provided as default mappings in the repository, implementing such mappings requires additions to the Max/MSP pipeline, since the values are already acquired in the current Max implementations. Therefore, we treat the slider as a ready sensor for real-time parameter modulation that can be tailored to specific experimental designs.

### 4.2 Social configuration: solo vs. dual use

One key aspect of the eMB Roma is that it supports both solo and dual modes. For convenience, the repository includes two ready-to-use Max/MSP patches: one configured for solo use and one for dual use. Although it is technically possible to run more than two eMBs from a single computer, we do not include code for multi-user setups beyond dyads because such configurations were not yet used or validated in our experiments.

Musical complexity configurations apply to both social modes, but dual mode expands the number of possible combinations. Figure 6 illustrates three representative dual use cases. In a “unison duet”, both eMBs play the same instrument and identical musical sequence; in a “counterpoint duet”, the devices play different but complementary musical lines (they may use the same or different timbres); and in a “band-like duet” one eMB can produce primarily melodic polyphonic streams (e.g., saxophone and keyboards) while the other provides rhythmic and harmonic accompaniment (e.g., drums and bass). This case example of band-like arrangement is shown in the example data in Figure 5, where each eMB polyphonic instrumental stream is visible in the MIDI data (panel A).

Beyond musical content, dual setups also permit configuration of inter-device relations such as the spatial alignment of rotation (aligned vs. mirrored) and the starting angular reference (the rotation angle defined as 0°). The starting angle determines where each eMB’s rotation is considered to begin, and it is set in the device firmware, as well as the direction of rotation (clockwise vs. anticlockwise), as explained in the section 2.3. For side-by-side participants, identical starting angles and rotating directions are typically simpler and often preferable. Nevertheless, for face-to-face dyads, initializing the two eMBs at opposite starting angles (12 o’clock and 6 o’clock) while both rotate clockwise produces mirrored movements that seem to facilitate interpersonal synchrony as both participants produce more spatially congruent movements.

## 5 Discussion

In this section, we discuss the technical and conceptual contributions of the e-Music Box Roma as an open and reproducible tool for music-based behavioral and neurophysiological research. We first outline the system’s main design features and introduce musical complexity and social configuration as two core dimensions for structuring experimental paradigms with the device. We then discuss potential applications across research, educational, clinical, and therapeutic contexts, and consider how the eMB Roma contributes to broader methodological and conceptual approaches to studying music, coordination, and social interaction. Finally, we address the practical accessibility and openness of the platform, discuss current limitations and future development directions, and conclude with an overview of its potential as a flexible tool for interdisciplinary research and translational applications.

The e-Music Box Roma offers a reproducible platform for music-based behavioral and neurophysiological research by combining a flexible open-source hardware and software environment. Its modular 3D-printed design and openly available firmware and software make it accessible, while its high temporal precision and MIDI compatibility support a wide range of experimental paradigms, ranging from simple solo timing tasks to complex, polyphonic joint performances, configured by varying musical complexity and social configuration parameters. An operator-facing Max/MSP GUI serves as the experiment’s central control hub and includes a TTL-trigger module that further enhances its integration with external systems, simplifying multimodal data acquisition paradigms, which are becoming more relevant in the behavioral research field (De Felice et al., 2025; Maidhof et al., 2014; Ohayon & Gordon, 2025; Varni et al., 2010).

The two configuration dimensions, musical complexity and social configuration, provide a general framework for designing and comparing experimental paradigms with the eMB Roma. Varying musical complexity (from single-line, monophonic material to multi-stream, polyphonic arrangements and different stimulus relationships) lets operators probe how task demands on attention, prediction, and sensorimotor coordination change with stimulus richness; varying social configuration (solo vs. dyad, with aligned or mirrored rotational mappings) isolates the contribution of interpersonal information and spatial alignment to coordination strategies. Together these axes make it straightforward to construct targeted contrasts (for example, unison versus counterpoint duets to test entrainment mechanisms, or solo versus band-like multi-stream tasks to examine multitasking and adaptive timing) while keeping other factors constant. Importantly, hardware-grounded settings such as starting angular reference, rotation direction, and perceptual access can be layered onto these dimensions to create paradigms that emphasize visual prediction, spatial coupling, or sensorimotor remapping; we provide recommended configuration files and firmware notes in Methods to support reproducible implementation. Finally, because the same core mappings and logging conventions are used across conditions, the dimensions facilitate cross-study comparisons and incremental extensions (e.g., larger groups or alternative meters) without changing the fundamental interaction logic, making it easier to link behavioral, physiological, and computational measures to specific manipulations.

Importantly, the eMB Roma expands access to active music making beyond trained musicians. Its intuitive control interface could allow näıve participants, including children, older adults, and clinical populations, to engage in musical interaction. This opens promising research and translational applications, including rehabilitation, developmental studies, and music therapy (Altenmüller & Schlaug, 2015; Backer & Sutton, 2014; Braun Janzen et al., 2022; Caputo et al., 2025; Fachner & Yap, 2025; Fachner et al., 2023; Farina et al., 2025; Hackney & Earhart, 2010; Kamp et al., 2025; MacRitchie et al., 2024; Thaut & McIntosh, 2014; Ting et al., 2015). For example, patients with motor or cognitive impairments could benefit from tasks that are both engaging, intrinsically rewarding (Zatorre, 2015) and precisely measurable, allowing operators to monitor progress over time. Similarly, studies with infants or young children could use the eMB Roma to examine the early emergence of sensorimotor synchronization and social responsiveness through naturalistic, music-based paradigms, domains that may be difficult to investigate systematically with traditional instruments.

Beyond its technical implementation, the eMB Roma contributes conceptually to the broader use of music as a platform for investigating human synchronization, social interaction, communication, and sensorimotor learning. Music-making provides an ecologically valid yet experimentally controllable framework in which individuals generate temporally structured actions, respond adaptively to auditory feedback, and synchronize with others (D’Ausilio et al., 2015). The eMB Roma translates rotary hand movements into musical output while simultaneously acquiring quantitative movement and performance data, pro-viding a reproducible, musically meaningful research platform. In this way, it bridges artistic and scientific perspectives, extending the methodological framework established by the original eMB (Novembre et al., 2015) and expanding it toward new domains.

So, how open and accessible is the eMB Roma in practice? To obtain a fully functioning device, operators need first to 3D-print the mechanical components, purchase the listed off-the-shelf electronic parts, and assemble the system following the provided technical documentation. Then, once the eMB is built and connected to a computer (using the available firmware code), it can be operated by downloading Max 8 along with the open-source patch. After updating the configuration files to match local specifications (e.g., COM ports of the eMB or paths), the device is operational. Importantly, no paid software license is required to run the provided patch; a Cycling ‘74 (Max/MSP 8) license is only necessary for those who wish to modify or extend the patch. This ensures that the eMB Roma can be adopted in both research and educational settings without prohibitive technical or financial barriers. Despite these advantages, the current implementation presents some limitations. MIDI transmission latency may also constrain temporal resolution, which is a limitation that affects many popular settings (Schultz, 2019). Moreover, wired serial communication may introduce additional latency and limit users mobility during the performances, which could be addressed in future developments by integrating wireless data transmission protocols such as Lab Streaming Layer (LSL). Future developments will aim to create a fully open-source software environment independent of Max/MSP (e.g., PureData), enabling free modification of the eMB software beyond read-only access, further expanding flexibility and accessibility. In summary, the eMB Roma represents a significant step toward open, reproducible, versatile, and widely accessible digital musical instruments for behavioral science, while providing controlled measures of the user’s movements and musical outputs. By combining a precise open-science experimental tool with expressive qualities of musical production, the eMB Roma provides a robust platform for investigating interpersonal synchronization, learning, therapy, and social behavior. At the same time, it supports both experimental control and creative exploration. The convergence of artistic expression, behavioral measurement, and clinical applicability of the eMB Roma highlights the potential of open digital musical instruments to bridge basic and applied research.

## Declarations

## Acknowledgments

We thank Flavia Arnese for her help with revising the manuscript and repository materials. Some icons used in the figures were adapted from The Noun Project (https://thenounproject. com) under CC BY 3.0 license. These include icons by Sherrinford, Firza Alamsyah, Uswa KDT, Lewin Design, Pentagon88, Creative Icon, H Alberto Gongora, Vectors Market, Gruffy Studio, At Her Cozy Corner, Kalaakarini, and mumur.

## Ethics approval

Not applicable.

## Consent to participate

Not applicable.

## Consent for publication

Not applicable.

## Availability of data and materials

The e-Music Box Roma is fully open-source. Hardware designs, firmware, and software are available at: https://github.com/sarafernandezab/eMusic Box Roma. The system can be assembled using off-the-shelf components and 3D-printed parts. The musical materials used in this study (original MIDI files and MIDI-derived songs eMB-ready) are available in the online repository.

## Code availability

All source code associated with the e-Music Box Roma is available at: https://github.com/ sarafernandezab/eMusic Box Roma.

The repository includes firmware for the embedded eMB devices, Max/MSP patches for experimental control, and Python scripts for generating stimulus from MIDI, generating configuration files, and loading data logging.

## Authors’ contributions

S.F.A. led the project and was responsible for conceptualization, implementation, and manuscript preparation. F.B. contributed to project development and manuscript revision. G.N. supervised the project and contributed to conceptualization and manuscript revision. L.O. and A.P. contributed to the conceptualization and mechanical design of the system. C.L. and M.C. contributed to the electronic design and assembly. P.E.K. provided feedback on the final manuscript.

## Supplementary Materials

**Table S1:**
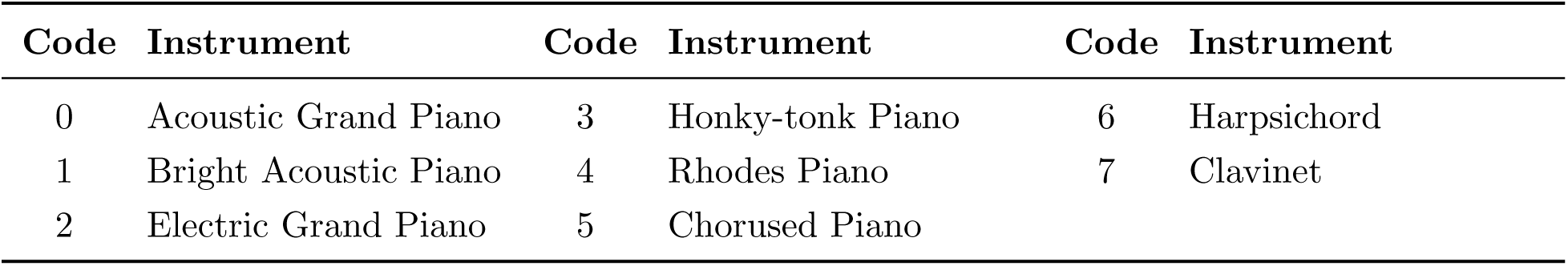
General MIDI piano instruments.

| Code | Instrument | Code | Instrument | Code | Instrument |
| --- | --- | --- | --- | --- | --- |
| 0 | Acoustic Grand Piano | 3 | Honky-tonk Piano | 6 | Harpsichord |
| 1 | Bright Acoustic Piano | 4 | Rhodes Piano | 7 | Clavinet |
| 2 | Electric Grand Piano | 5 | Chorused Piano |  |  |

**Table S2:** General MIDI chromatic percussion instruments.

| Code | Instrument | Code | Instrument | Code | Instrument |
| --- | --- | --- | --- | --- | --- |
| 8 | Celesta | 11 | Vibraphone | 14 | Tubular Bells |
| 9 | Glockenspiel | 12 | Marimba | 15 | Dulcimer |
| 10 | Music Box | 13 | Xylophone |  |  |

**Table S3:** General MIDI organ instruments.

| Code | Instrument | Code | Instrument | Code | Instrument |
| --- | --- | --- | --- | --- | --- |
| 16 | Hammond Organ | 19 | Church Organ | 22 | Harmonica |
| 17 | Percussive Organ | 20 | Reed Organ | 23 | Tango Accordion |
| 18 | Rock Organ | 21 | Accordion |  |  |

**Table S4:** General MIDI guitar instruments.

| Code | Instrument | Code | Instrument | Code | Instrument |
| --- | --- | --- | --- | --- | --- |
| 24 | Acoustic Nylon Guitar | 27 | Electric Clean Guitar | 30 | Distortion Guitar |
| 25 | Acoustic Steel Guitar | 28 | Electric Muted Guitar | 31 | Guitar Harmonics |
| 26 | Electric Jazz Guitar | 29 | Overdriven Guitar |  |  |

**Table S5:** General MIDI bass instruments.

| Code | Instrument | Code | Instrument | Code | Instrument |
| --- | --- | --- | --- | --- | --- |
| 32 | Acoustic Bass | 35 | Fretless Bass | 38 | Synth Bass 1 |
| 33 | Fingered Electric Bass | 36 | Slap Bass 1 | 39 | Synth Bass 2 |
| 34 | Plucked Electric Bass | 37 | Slap Bass 2 |  |  |

**Table S6:** General MIDI string instruments.

| Code | Instrument | Code | Instrument | Code | Instrument |
| --- | --- | --- | --- | --- | --- |
| 40 | Violin | 43 | Contrabass | 46 | Orchestral Harp |
| 41 | Viola | 44 | Tremolo Strings | 47 | Timpani |
| 42 | Cello | 45 | Pizzicato Strings |  |  |

**Table S7:** General MIDI ensemble instruments.

| Code | Instrument | Code | Instrument | Code | Instrument |
| --- | --- | --- | --- | --- | --- |
| 48 | String Ensemble 1 | 51 | Synth Strings 2 | 54 | Synth Voice |
| 49 | String Ensemble 2 | 52 | Choir Aah | 55 | Orchestral Hit |
| 50 | Synth Strings 1 | 53 | Choir Ooh |  |  |

**Table S8:** General MIDI brass instruments.

| Code | Instrument | Code | Instrument | Code | Instrument |
| --- | --- | --- | --- | --- | --- |
| 56 | Trumpet | 59 | Muted Trumpet | 62 | Synth Brass 1 |
| 57 | Trombone | 60 | French Horn | 63 | Synth Brass 2 |
| 58 | Tuba | 61 | Brass Section |  |  |

**Table S9:** General MIDI reed instruments.

| Code | Instrument | Code | Instrument | Code | Instrument |
| --- | --- | --- | --- | --- | --- |
| 64 | Soprano Sax | 67 | Baritone Sax | 70 | Bassoon |
| 65 | Alto Sax | 68 | Oboe | 71 | Clarinet |
| 66 | Tenor Sax | 69 | English Horn |  |  |

**Table S10:** General MIDI pipe instruments.

| Code | Instrument | Code | Instrument | Code | Instrument |
| --- | --- | --- | --- | --- | --- |
| 72 | Piccolo | 75 | Pan Flute | 78 | Whistle |
| 73 | Flute | 76 | Bottle Blow | 79 | Ocarina |
| 74 | Recorder | 77 | Shakuhachi |  |  |

**Table S11:** General MIDI percussion (Channel 10)

| Note | Instrument | Note | Instrument | Note | Instrument |
| --- | --- | --- | --- | --- | --- |
| 35 | Acoustic Bass Drum | 51 | Ride Cymbal 1 | 67 | High Agogo |
| 36 | Bass Drum 1 | 52 | Chinese Cymbal | 68 | Low Agogo |
| 37 | Side Stick | 53 | Ride Bell | 69 | Cabasa |
| 38 | Acoustic Snare | 54 | Tambourine | 70 | Maracas |
| 39 | Hand Clap | 55 | Splash Cymbal | 71 | Short Whistle |
| 40 | Electric Snare | 56 | Cowbell | 72 | Long Whistle |
| 41 | Low Floor Tom | 57 | Crash Cymbal 2 | 73 | Short Guiro |
| 42 | Closed Hi-Hat | 58 | Vibraslap | 74 | Long Guiro |
| 43 | High Floor Tom | 59 | Ride Cymbal 2 | 75 | Claves |
| 44 | Pedal Hi-Hat | 60 | High Bongo | 76 | High Wood Block |
| 45 | Low Tom | 61 | Low Bongo | 77 | Low Wood Block |
| 46 | Open Hi-Hat | 62 | Mute High Conga | 78 | Mute Cuica |
| 47 | Low Mid Tom | 63 | Open High Conga | 79 | Open Cuica |
| 48 | High Mid Tom | 64 | Low Conga | 80 | Mute Triangle |
| 49 | Crash Cymbal 1 | 65 | High Timbale | 81 | Open Triangle |
| 50 | High Tom | 66 | Low Timbale |  |  |

## References

Abalde, S. F., Rigby, A., Keller, P. E., & Novembre, G. (2024). A framework for joint music making: Behavioral findings, neural processes, and computational models. Neuroscience & Biobehavioral Reviews, 167, 105816. 10.1016/j.neubiorev.2024.105816

Altenmüller, E., & Schlaug, G. (2015). Apollo’s gift: New aspects of neurologic music therapy. Prog Brain Res, 217, 237–52.

Backer, J. D., & Sutton, J. (2014). The Music in Music Therapy: Psychodynamic Music Therapy in Europe: Clinical, Theoretical and Research Approaches. Jessica Kingsley Publishers.

Bigand, F., Bianco, R., Abalde, S. F., Nguyen, T., & Novembre, G. (2025). EEG of the Dancing Brain: Decoding Sensory, Motor, and Social Processes during Dyadic Dance. Journal of Neuroscience, 45 (21). 10.1523/JNEUROSCI.2372-24.2025

Bigand, F., Bianco, R., Abalde, S. F., & Novembre, G. (2024). The geometry of interpersonal synchrony in human dance. Current Biology, 34 (13), 3011–3019.e4. 10.1016/j.cub.2024.05.055

Braun Janzen, T., Koshimori, Y., Richard, N. M., & Thaut, M. H. (2022). Rhythm and Music-Based Interventions in Motor Rehabilitation: Current Evidence and Future Perspectives. Frontiers in Human Neuroscience, 15. 10.3389/fnhum.2021.789467

Caputo, C. C., Pranjíc, M., Koshimori, Y., & Thaut, M. H. (2025). The Effects of Music-Based Patterned Sensory Enhancement on Motor Function: A Scoping Review. Brain Sciences, 15 (7), 664. 10.3390/brainsci15070664

Coorevits, E., Maes, P.-J., Six, J., & Leman, M. (2020). The influence of performing gesture type on interpersonal musical timing, and the role of visual contact and tempo. Acta Psychologica, 210, 103166. 10.1016/j.actpsy.2020.103166

D’Ausilio, A., Novembre, G., Fadiga, L., & Keller, P. E. (2015). What can music tell us about social interaction? Trends in Cognitive Sciences, 19 (3), 111–114. 10.1016/j.tics.2015.01.005

De Felice, S., Chand, T., Croy, I., Engert, V., Goldstein, P., Holroyd, C. B., Kirsch, P., Krach, S., Ma, Y., Scheele, D., Schurz, M., Schweinberger, S. R., Hoehl, S., & Vrticka, P. (2025). Relational neuroscience: Insights from hyperscanning research. Neuroscience & Biobehavioral Reviews, 169, 105979. 10.1016/j.neubiorev.2024.105979

Duarte, E. G., Cossette, I., & Wanderley, M. M. (2023). Analysis of Accessible Digital Musical Instruments through the lens of disability models: A case study with instruments targetingd/Deaf people. Frontiers in Computer Science, 5, 1158476. 10.3389/fcomp.2023.1158476

Fachner, J., Maidhof, C., Murtagh, D., De Silva, D., Pasqualitto, F., Fernie, P., Panin, F., Michell, A., Muller-Rodriguez, L., & Odell-Miller, H. (2023). Music therapy, neural processing, and craving reduction: An RCT protocol for a mixed methods feasibility study in a Community Substance Misuse Treatment Service. Addiction Science & Clinical Practice, 18 (1), 36. 10.1186/s13722-023-00385-y

Fachner, J., & Yap, S. S. (2025). Make the Invisible Visible - Towards Process-Based Outcome Research on Mechanisms of Change in Music Therapy. In A. Raglio (Ed.), Music and Music Therapy Interventions in Clinical Practice: A Textbook for Music Therapy Professionals, Clinicians and Researchers (pp. 471–488). Springer Nature Switzerland. 10.1007/978-3-031-88578-520

Farina, N., Brewster, S., Vito, P. D. C. S., Fachner, J., Street, A., Miranda, E., & Banerjee, S. (2025). Real-time detection of agitation in people with dementia: The RadioMe system. Alzheimer’s & Dementia, 21 (S3), e099249. 10.1002/alz70857 099249

Feng, Y.-L., Wang, Z., Yao, Y., Li, H., Diao, Y., Peng, Y., & Mi, H. (2024). Co-designing the Collaborative Digital Musical Instruments for Group Music Therapy. Proceedings of the CHI Conference on Human Factors in Computing Systems, 1–18. 10.1145/3613904.3642649

Fink, L., Alexander, P., & Janata, P. (2022). The Groove Enhancement Machine (GEM): A Multi-Person Adaptive Metronome to Manipulate Sensorimotor Synchronization and Subjective Enjoyment. Frontiers in Human Neuroscience, 16, 916551. 10.3389/fnhum.2022.916551

Hackney, M. E., & Earhart, G. M. (2010). Effects of Dance on Gait and Balance in Parkinson’s Disease: A Comparison of Partnered and Nonpartnered Dance Movement. Neurorehabilitation and Neural Repair, 24 (4), 384–392. 10.1177/1545968309353329

Jack, R., Harrison, J., & McPherson, A. (2020). Digital Musical Instruments as Research Products. 10.5281/ZENODO.4813465

Kamp, D., Bolis, D., & Schilbach, L. (2025). Understanding and explaining differences across minds in social interaction: Insights from social neuroscience and clinical psychiatry. European Archives of Psychiatry and Clinical Neuroscience, 275 (8), 2199–2201. 10.1007/s00406-025-02136-3

Kavety, S. (2019). Digital music making: Developing a method for using technology in music psychotherapy. https://api.semanticscholar.org/CorpusID:182271686

Keller, P. E., & Rieger, M. (2009). Special Issue—Musical Movement and Synchronization. Music Perception: An Interdisciplinary Journal, 26 (5), 397–400. 10.1525/mp.2009.26.5.397

Liebermann-Jordanidis, H., Novembre, G., Koch, I., & Keller, P. E. (2021). Simultaneous self-other integration and segregation support real-time interpersonal coordination in a musical joint action task. Acta Psychologica, 218, 103348. 10.1016/j.actpsy.2021.103348

Lindetorp, H., Svahn, M., Hölling, J., Falkenberg, K., & Frid, E. (2023). Collaborative music-making: Special educational needs school assistants as facilitators in performances with accessible digital musical instruments. Frontiers in Computer Science, 5, 1165442. 10.3389/fcomp.2023.1165442

MacRitchie, J., Breaden, M., Taylor, J. R., & Milne, A. J. (2024). Exploring older adult needs and preferences for technology-assisted group music-making. A qualitative analysis of data collected during the participatory user-centred design process. Disability and Rehabilitation: Assistive Technology, 19 (5), 1935–1944. 10.1080/17483107.2022.2077461

Maes, P.-J., Kerrebroeck, B. van, Rosso, M., Marouda, I., & Leman, M. (2024). Extended reality (XR) in embodied musical art and science. In The Routledge Handbook of Embodied Cognition (2nd ed.). Routledge.

Maidhof, C., Kästner, T., & Makkonen, T. (2014). Combining EEG, MIDI, and motion capture techniques for investigating musical performance. Behavior Research Methods, 46 (1), 185–195. 10.3758/s13428-013-0363-9

MIDI Manufacturers Association. (1996). Complete midi 1.0 detailed specification [Document Version 4.2]. MIDI Manufacturers Association.

Miranda, E. R., & Wanderley, M. M. (2006). New Digital Musical Instruments: Control and Interaction Beyond the Keyboard. A-R Editions, Inc.

Nicol, J., Loehr, J., Christensen, J., Lang, J., & Peacock, S. (2024). Duet playing in dementia care: A new therapeutic music technology. Disability and Rehabilitation: Assistive Technology, 19 (8), 3139–3152. 10.1080/17483107.2024.2351498

Novembre, G., Mitsopoulos, Z., & Keller, P. E. (2019). Empathic perspective taking promotes interpersonal coordination through music. Scientific Reports, 9 (1), 12255. 10.1038/s41598-019-48556-9

Novembre, G., Varlet, M., Muawiyath, S., Stevens, C. J., & Keller, P. E. (2015). The E-music box: An empirical method for exploring the universal capacity for musical production and for social interaction through music. Royal Society Open Science, 2 (11), 150286. 10.1098/rsos.150286

Ohayon, S., & Gordon, I. (2025). Multimodal interpersonal synchrony: Systematic review and meta-analysis. Behavioural Brain Research, 480, 115369. 10.1016/j.bbr.2024.115369

Partesotti, E., Feitosa, J. A., Manzolli, J., & Castellano, G. (2025). Stimulating neuroplasticity: Therapeutic applications of an extended digital musical instrument. Nordic Journal of Music Therapy, 34 (1), 62–80. 10.1080/08098131.2024.2445824

Pikovsky, A., Rosenblum, M., & Kurths, J. (2001, October). Synchronization: A Universal Concept in Nonlinear Sciences. Cambridge University Press.

Raffel, C., & Ellis, D. P. W. (2014). Intuitive analysis, creation and manipulation of midi data with pretty*_m_idi*. Proceedings of the 15th International Conference on Music Information Retrieval (ISMIR) Late Breaking and Demo Papers.

Repp, B. H., & Su, Y.-H. (2013). Sensorimotor synchronization: A review of recent research (2006-2012). Psychonomic Bulletin & Review, 20 (3), 403–452. 10.3758/s13423-012-0371-2

Rosenbaum, D. A. (2005). The Cinderella of Psychology: The Neglect of Motor Control in the Science of Mental Life and Behavior. American Psychologist, 60 (4), 308–317. 10.1037/0003-066X.60.4.308

Savage, P. E., Brown, S., Sakai, E., & Currie, T. E. (2015). Statistical universals reveal the structures and functions of human music. Proceedings of the National Academy of Sciences, 112 (29), 8987–8992. 10.1073/pnas.1414495112

Schultz, B. G. (2019). The Schultz MIDI Benchmarking Toolbox for MIDI interfaces, percussion pads, and sound cards. Behavior Research Methods, 51 (1), 204–234. 10.3758/s13428-018-1042-7

Sihvonen, A. J., Särkämö, T., Leo, V., Tervaniemi, M., Altenmüller, E., & Soinila, S. (2017). Music-based interventions in neurological rehabilitation. The Lancet Neurology, 16 (8), 648–660. 10.1016/S1474-4422(17)30168-0

Thaut, M. H., & McIntosh, G. C. (2014). Neurologic Music Therapy in Stroke Rehabilitation. Current Physical Medicine and Rehabilitation Reports, 2 (2), 106–113. 10.1007/s40141-014-0049-y

Ting, L. H., Chiel, H. J., Trumbower, R. D., Allen, J. L., McKay, J. L., Hackney, M. E., & Kesar, T. M. (2015). Neuromechanical Principles Underlying Movement Modularity and Their Implications for Rehabilitation. Neuron, 86 (1), 38–54. 10.1016/j.neuron.2015.02.042

Toiviainen, P., & Keller, P. E. (2010). Special Issue: Spatiotemporal Music Cognition. Music Perception, 28 (1), 1–1. 10.1525/mp.2010.28.1.1

Varni, G., Volpe, G., & Camurri, A. (2010). A System for Real-Time Multimodal Analysis of Nonverbal Affective Social Interaction in User-Centric Media. IEEE Transactions on Multimedia, 12 (6), 576–590. 10.1109/TMM.2010.2052592

Wanderley, M. M., & Depalle, P. (2004). Gestural control of sound synthesis. Proceedings of the IEEE, 92 (4), 632–644. 10.1109/JPROC.2004.825882

Zatorre, R. J. (2015). Musical pleasure and reward: Mechanisms and dysfunction. Annals of the New York Academy of Sciences, 1337 (1), 202–211. 10.1111/nyas.12677

Zayas-Garin, E., & McPherson, A. (2022). Dialogic Design of Accessible Digital Musical Instruments: Investigating Performer Experience [Conference Name: NIME 2022 Place: The University of Auckland, New Zealand]. NIME 2022. 10.21428/92fbeb44.2b8ce9a4

Zhou, Z., Christensen, J., Cummings, J. A., & Loehr, J. D. (2023). Not just in sync: Relations between partners’ actions influence the sense of joint agency during joint action. Consciousness and Cognition, 111, 103521. 10.1016/j.concog.2023.103521

